# Structural polymorphism of amyloid fibrils in cardiac transthyretin amyloidosis revealed by cryo-electron microscopy

**DOI:** 10.1101/2022.06.21.496949

**Authors:** Binh An Nguyen, Shumaila Afrin, Virender Singh, Yasmin Ahmed, Rose Pedretti, Maria del Carmen Fernandez-Ramirez, Merrill Douglas Benson, Michael Sawaya, Preeti Singh, Qin Cao, David Boyer, Alexander Pope, Pawel Wydorski, Siddharth Kumar, Farzeen Chhapra, David Eisenberg, Lorena Saelices

**Affiliations:** Center for Alzheimer’s and Neurodegenerative Diseases, Department of Biophysics, University of Texas Southwestern Medical Center (UTSW), Dallas, TX, USA; Department of Pathology and Laboratory Medicine, Indiana University School of Medicine, Indianapolis, IN, USA; Department of Biological Chemistry, University of California, Los Angeles, Howard Hughes Medical institute, CA, USA; Key Laboratory for the Genetics of Developmental and Neuropsychiatric Disorders (Ministry of Education), Bio-X Institutes, Shanghai Jiao Tong University, Shanghai, China; Department of Neuroscience, University of Texas Southwestern Medical Center, Dallas, TX, USA

**Author notes:** These authors contributed equally (Binh An Nguyen, Shumaila Afrin). Correspondence to: Lorena Saelices Gómez.

## Abstract

The deposition of amyloidogenic transthyretin (ATTR) in ATTR amyloidosis leads to an unexplained variety of clinical phenotypes, including cardiomyopathy. In brain amyloid conditions, there is an apparent association between the clinical phenotype and the amyloid fibril structure. Here, we question this phenotype-structure association in cardiac amyloidoses by determining the cryo-electron microscopy structures of fibrils extracted from the hearts of seven ATTR amyloidosis patients. We found that, in contrast to brain fibrils, cardiac ATTR fibrils display a structural polymorphism that is not genotype-specific, can co-exist within the same individual, and is independent of the cardiac phenotype. This polymorphism challenges the current paradigm of “one disease equals one fibril fold” proposed in tauopathies and synucleinopathies, and questions whether a similar structural heterogeneity occurs in other amyloidoses.

**One-Sentence Summary:** Unlike brain amyloid fibrils, cardiac ATTR fibrils are polymorphic independent of genotype and even within the same patient.

## Introduction

Amyloidoses are a group of fatal disorders caused by the pathological accumulation of amyloid fibrils in affected organs(*1*). In amyloid transthyretin (ATTR) amyloidosis, the amyloid fibrils are composed of the amyloidogenic form of the hormone transporter transthyretin(*2*). ATTR self-assemblage caused by mutations in the *TTR* gene leads to variant ATTR (ATTRv) amyloidosis. Other unknown processes related to age lead to wild-type ATTR (ATTRwt) amyloidosis. The clinical presentation of ATTRv amyloidosis is unpredictable and variable, often manifesting between 25 and 65 years of age with polyneuropathy, autonomic neuropathy, gastrointestinal and eye involvement, carpal tunnel syndrome, spinal canal stenosis, and/or cardiomyopathy(*3*). The clinical presentation of ATTRwt is more predictable and better characterized; it manifests later in life (>60 years old) as cardiomyopathy, and mainly affects men(*4*). The roots of the phenotypic variability of ATTR amyloidosis are yet unknown: is it connected to heterogeneity of fibril structures, or to characteristics of the surrounding tissue, or to other factors?

Recent cryo-electron microscopy (cryo-EM) studies on tauopathies and synucleinopathies, a set of brain amyloid diseases respectively associated with the deposition of tau and α-synuclein, have looked for the source of their poorly understood phenotypic variability(*5–8*). These studies reveal that fibrils from patients presenting the same disease share the same fold, or *polymorph*, supporting the idea that the fibril structure determines disease presentation(*9*). But studies focused on brain diseases offer limited information about whether amyloid formation is influenced by the type of tissue or mutation.

There are two studies that associate ATTR amyloidosis with the formation of distinct fibril polymorphs in two different organs: the heart(*10*) and the vitreous humor(*11*). Both cases involved the same pathological mutation, ATTRV30M; however, the small number of patients limits the studies’ strength and does not rule out whether mutations or tissue type could affect amyloid structure. In this study, we focus on the structure of cardiac ATTR fibrils to evaluate structural polymorphism in the heart. We used cryo-EM to determine the molecular structure of fibrils extracted from seven patients with ATTR amyloidosis - two ATTRwt cases and five ATTRv. We found that ATTR mutations perturb the fibril fold locally at the mutation site, suggesting that the degree of structural polymorphism in ATTR fibrils may be driven by the individual and their genotype. These results show that, contrary to tauopathies, ATTR amyloidosis patients can present structural variation in their fibrils, even when carrying the same variant. Additional studies on organ-specific structural heterogeneity and the influence of fibril folding in clinical presentations are warranted. A deeper structural understanding of ATTR amyloid fibrils could aid development of new therapeutic and diagnostic strategies to improve patients’ prognosis and quality of life.

## Results

### ATTRwt fibrils extracted from two different patients share a common core

We collected cryo-EM images of the fibrils extracted from the hearts of two ATTRwt amyloidosis patients, patient 1 and patient 2. Two-dimensional (2D) classification discerned two polymorphs that differ in their twisting (Figure 1A and Figure S3). The most prevalent species presented a clear twist, here referred to as *curvy* fibrils, and accounted for at least 95% of the extracted particles (Table S2). The *straight* species was not suitable for structure determination because it lacked a twist (Figure 1A).

**Figure 1.**
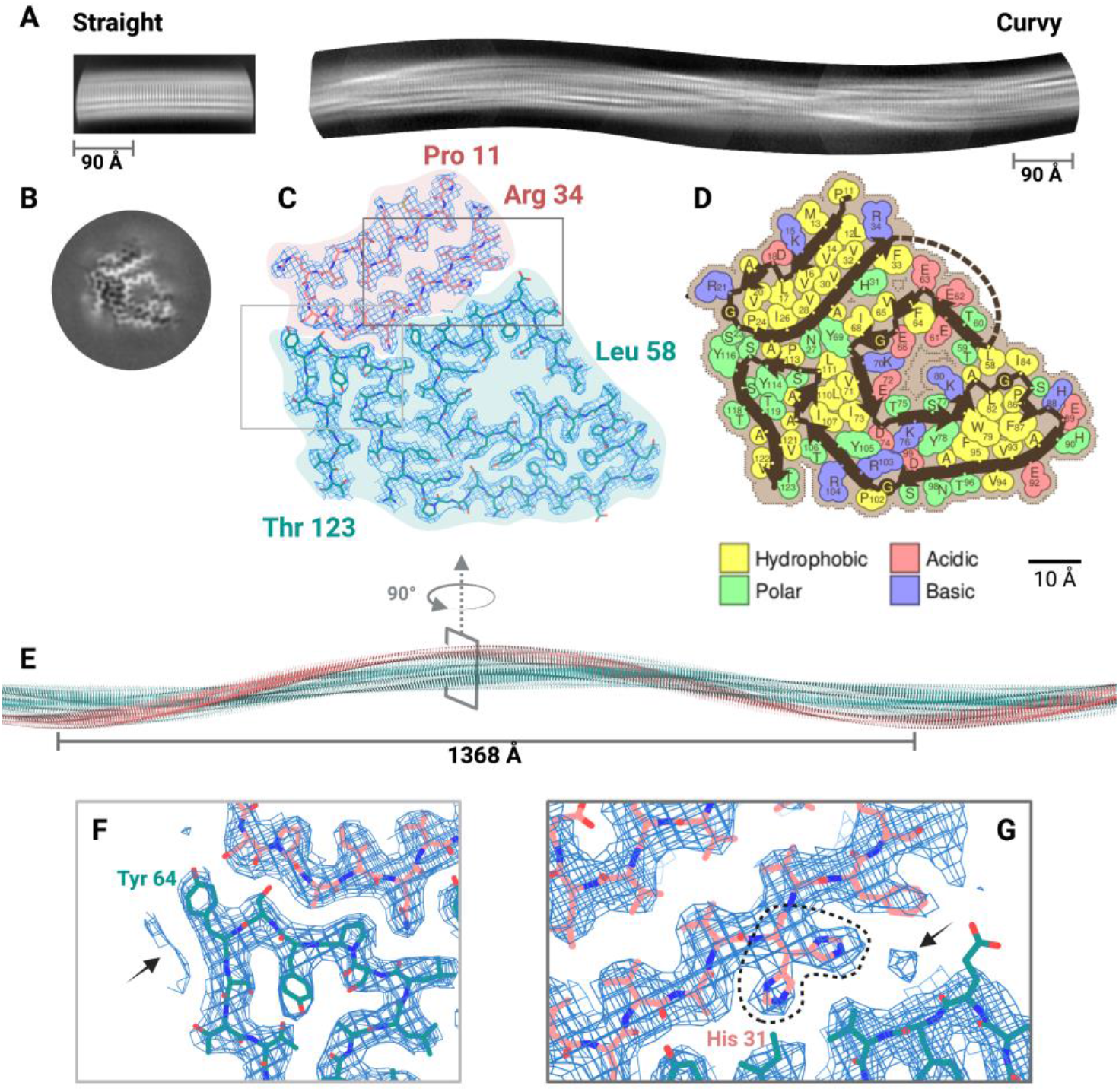
Cryo-EM structure of cardiac fibrils from a ATTRwt (A-G) patients showing a common conformation. **(A)**. Representative 2D class averages of straight fibrils (left) and curvy fibrils after stitching (right) from ATTRwt patient 1. **(B)**. Three-dimensional (3D) class averages of ATTRwt curvy fibrils. **(C)**. Cryo-EM density and atomic model of ATTRwt fibrils from patient 1. The model contains two fragments of transthyretin colored pink (residues Pro 11 to Arg 34) and turquoise (residues Leu 58 to Thr 123). **(D)**. Schematic view of the ATTRwt fibril core showing residue composition. **(E)**. Model reconstruction of the ATTRwt fibril. **(F)**. Close view of an unsatisfied density neighboring the residue Tyr 64 (marked with an arrow) in ATTRwt fibrils, apparent at a high contour level. **(G)**. Close view of the interface between the N and C terminal fragments in ATTRwt fibrils, pointing to unsatisfied densities apparent at a high contour level. One density blob could be assigned to an alternative rotamer of His 31 (dashed line).

Using helical reconstruction, we determined the cryo-EM structures of curvy ATTR fibrils extracted from the two patients. The overall resolution of the ATTRwt structures from patients 1 and 2 were 3.3 and 3.7 Å, respectively. Both structure cores shared the same fold (Figure 1B and Figure S3B), with one single protofilament composed of two ATTR fragments: Pro 11 to Lys 35 and Leu 58 to Thr 123 (Figure 1C). The overall secondary structure of each chain is summarized in Figure S5, and contains highly interdigitated interfaces mostly made of hydrophobic residues, also known as *steric zippers*(*12*), and a polar channel that involves residues from Leu 58 to Ile 84, which folds in a pentagon-like shape (Figure 1D). ATTRwt fibrils from patient 1 presented a crossover length of 684 Å with a helical twist of −1.25° and a helical rise of 4.75 Å (Figure 1E). ATTRwt fibrils from patient 2 presented a crossover length of 663 Å with a helical twist of −1.29° and a helical rise of 4.78 Å (Table S2).

### Curvy ATTRv fibrils come in different flavors

We extracted the fibrils from five ATTRv amyloidosis patients carrying the variants ATTRP24S, ATTRT60A, ATTRI84S, and ATTRV122I (Table S1). All five samples yielded a predominant number of curvy fibrils and a much lower number of straight fibrils as seen in ATTRwt fibrils during 2D classification (Table S2). The helical reconstruction of curvy fibrils from variants ATTRP24S and ATTRT60A yielded structures resembling those of ATTRwt fibrils in crossover distance, helical twist, helical rise, and overall structural fold (Figure S4 and Table S2). However, we observed structural differences in fibrils extracted from the other ATTRv genotypes.

Curvy fibrils extracted from the one ATTRV122I patient yielded two distinct curvy populations with different twists (Figure 2 and Table S2). These two populations share a similar structure fold, and identical refined helical rise (Figure 2C-F and Table S2). Additionally to the fibril twist, the two populations differ in that ATTRV122Ib fibrils contain two extended interfaces consisting of 4 additional residues at the beginning of the C-terminal segment (Figure 2E) and 4 additional residues at the end of the C-terminal segment (Figure 2F). The residue composition of these fibrils are shown in Figure S6.

**Figure 2.**
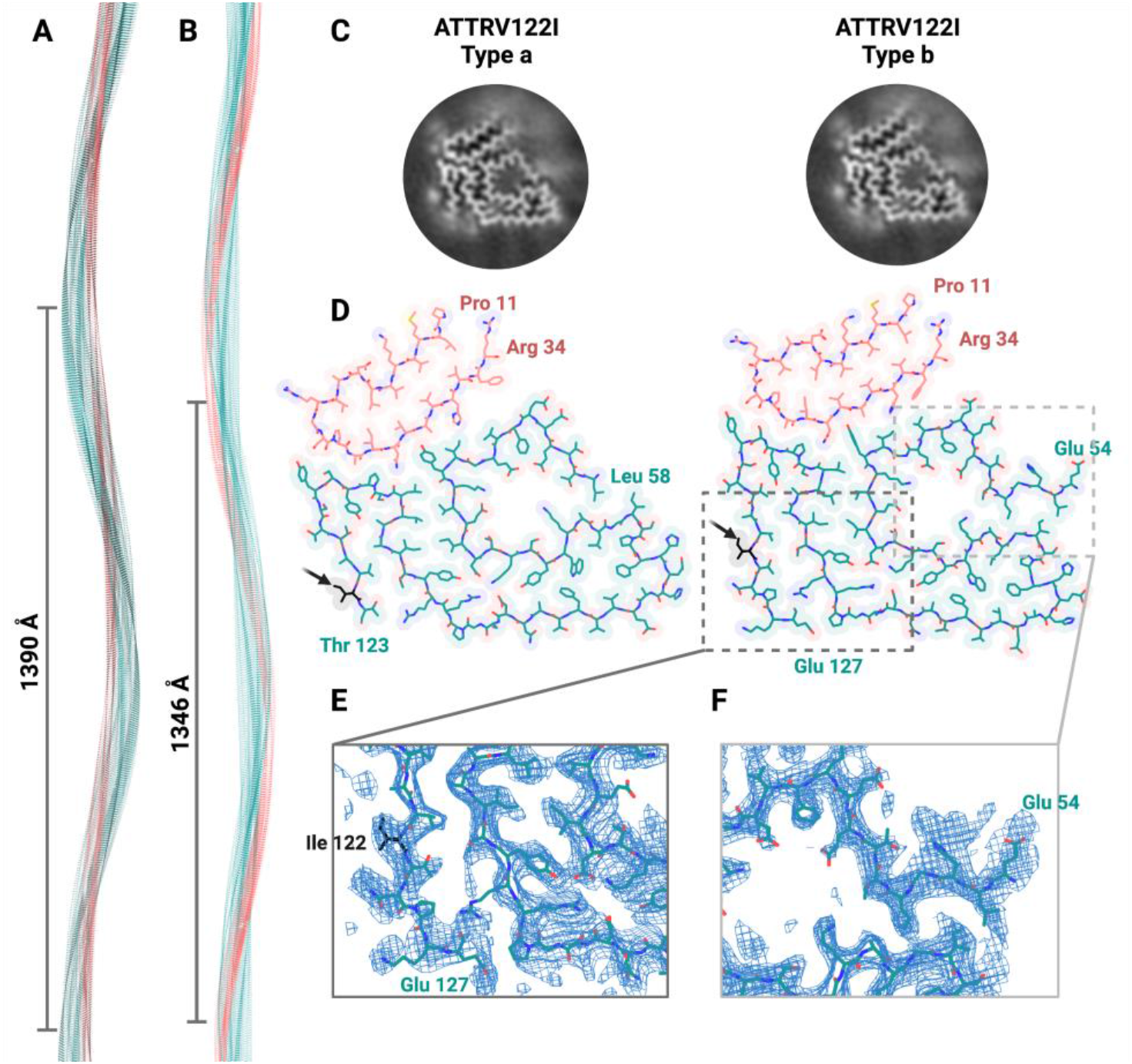
Cryo-EM structures of two distinct ATTRV122I fibrils extracted from the same patient. **(A and B)**. Model reconstructions of two distinct ATTRV122I fibrils from the same patient. They differ in the helical twist angle and crossover distance. **(C)**. 3D class averages of the two types of fibrils. **(D)**. Atomic model of ATTRV122I fibrils (Left, ATTRV122Ia; Right, ATTRV122Ib). The two fragments of transthyretin are colored pink (N terminus) and turquoise (C terminus). Arrows point at the mutation site. **(E and F)**. Enlarged view of the cryo-EM density and atomic model of the additional residues found in ATTRV122Ib fibrils: residues Glu 54 onwards (**E**) and Glu 127 backwards (**F**).

Curvy fibrils extracted from two ATTRI84S patients display structural differences (Figure 3). ATTRI84S fibrils from patient 1 lack the segment prior to Gly 67 of the C-terminal fragment, suggesting that this segment may be disordered or absent (Figure 3A-D). In contrast, the fibrils extracted from an additional ATTRI84S patient, patient 2 (Figure 3E-H), have a defined density for residues Thr 59 to Gly 67 (Figure 3F-H), one residue short to fibrils from ATTRwt or the other ATTRv patients.

**Figure 3.**
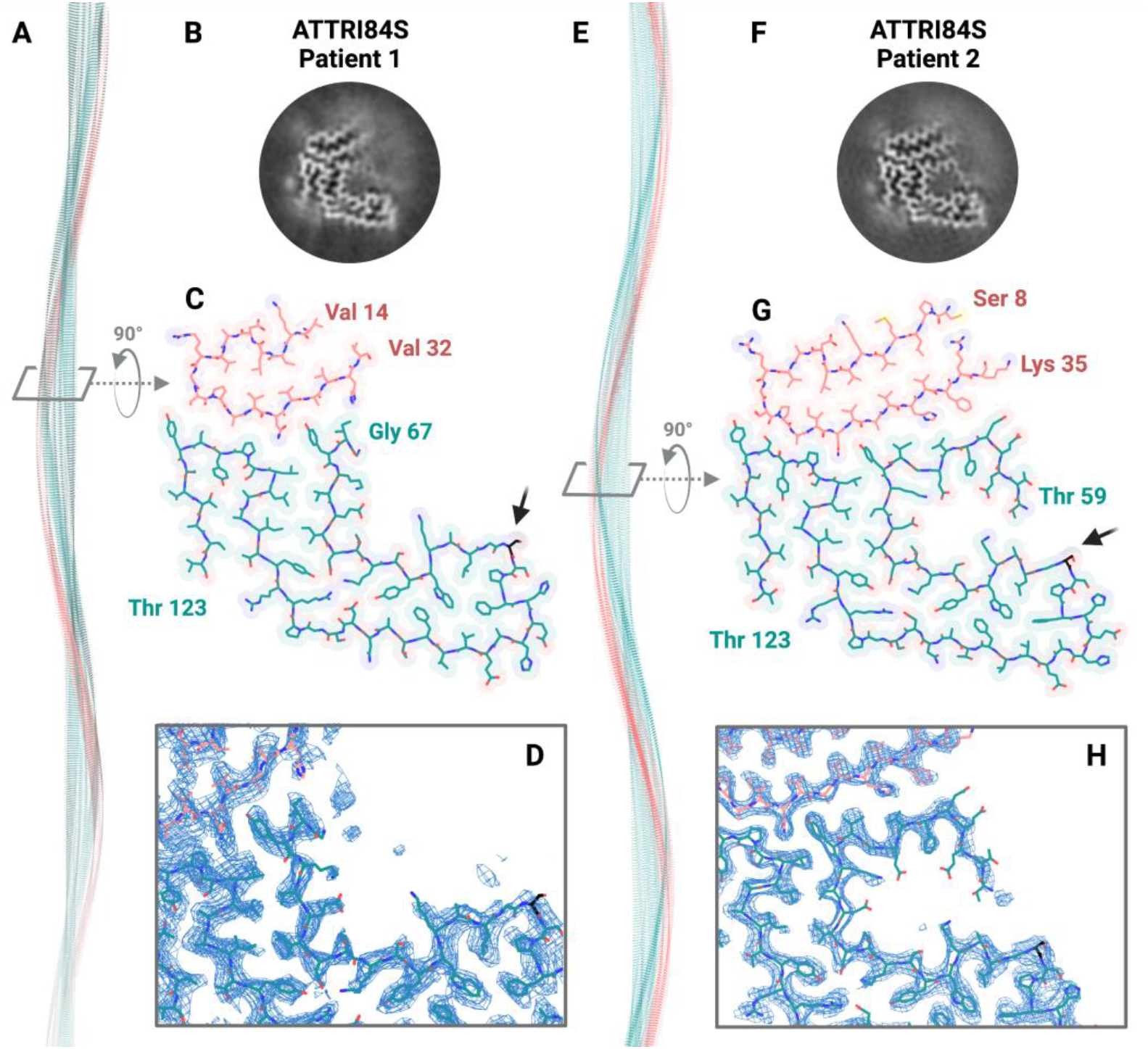
Cryo-EM structures of ATTRI84S fibrils from two patients, ATTRI84S patient 1 (A-D) and ATTRI84S patient 2 (E-H). **(A and E)**. Model reconstructions. The N terminal fragment of transthyretin is colored pink, and the C terminal is colored turquoise. (**B and F)**. 3D class averages. (**C and G)**. Atomic model of ATTRI84S fibrils, as labeled. The N terminal fragment of transthyretin is colored pink, and the C terminal is colored turquoise. Arrows point at the mutation site. (**D and H)**. Close view of the cryo-EM density and atomic model of the polar channel in ATTRI84S fibrils.

### ATTRv fibrils are associated with local structure perturbations

Local resolution maps of ATTR structures reveal additional differences between ATTRwt and ATTRv fibrils. Mutations appear to disturb the local structures at and near the mutation sites of the density maps (Figure 4A). In ATTRP24S fibrils and ATTRI84S fibrils from patient 1, the resolution near the mutation sites reaches ~6.5 Å and the overall map resolutions are 3.65 and 3.61 Å, respectively. In ATTRI84S fibrils from patient 2, the resolution near the mutation site reaches ~3.8 Å, and the overall map resolution is 3.07 Å. In ATTRV122I and ATTRT60A fibrils, the structural perturbation affects the surface of the fibril rather than the mutation site only. The resolution at the surface of the two types of ATTRV122I fibrils reaches ~4.2 Å, and the overall resolution of their maps is 3.16 and 3.2 Å. In ATTRT60A fibrils, the resolution at the surface reaches ~3.8 Å, and the overall resolution is 3.3 Å. In contrast, the resolution map of both ATTRwt fibrils is more homogeneous (Figure 4B).

**Figure 4.**
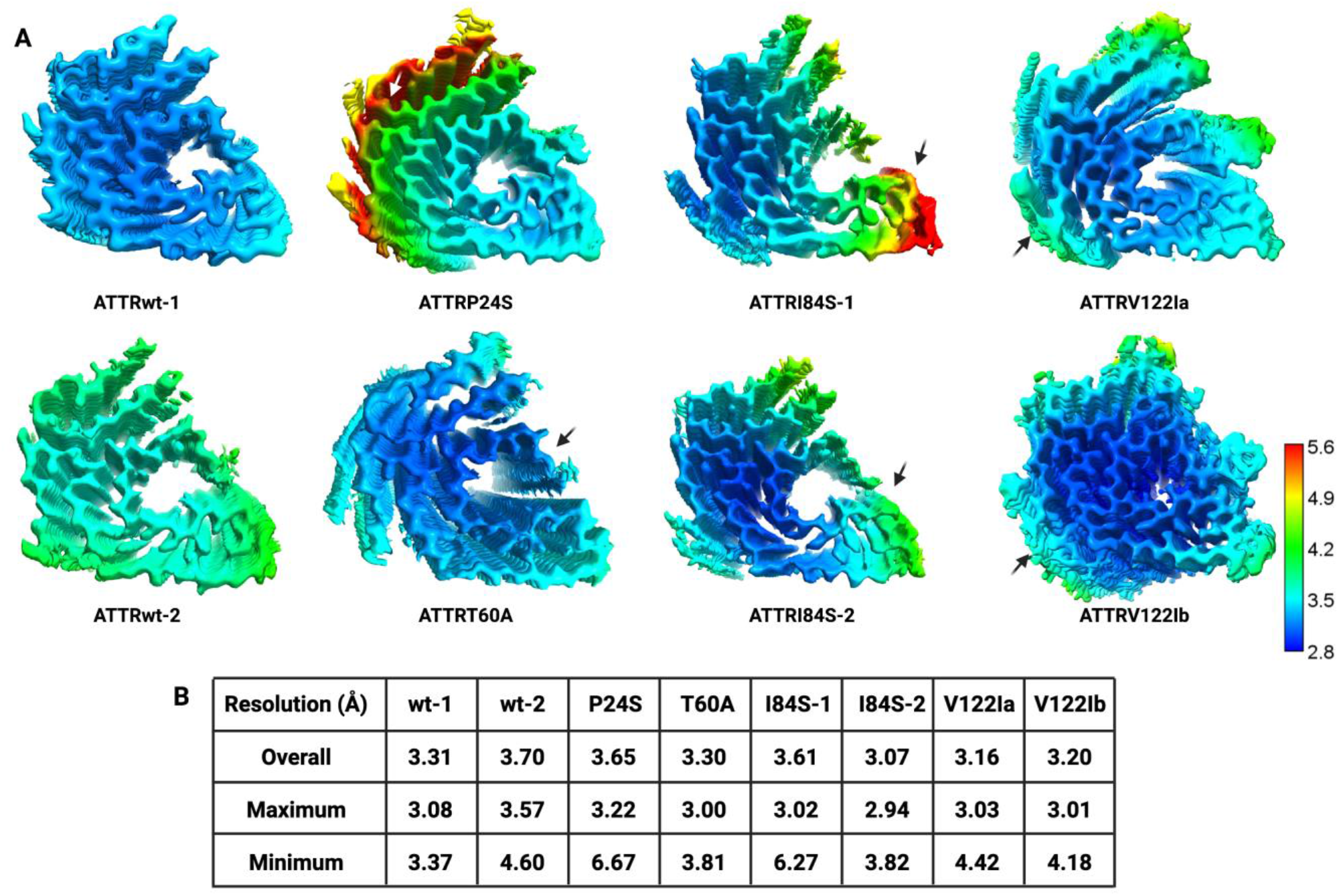
Local resolution maps of ATTR fibrils. **(A)**. Local resolution estimation for 3D reconstructions of ATTR fibrils. Arrows point to the locations of mutations on each density map. ATTRP24S mutation is marked with a white arrow to facilitate color contrast. Resolution scale in Angstroms. **(B)**. Summary of overall, maximum, and minimum resolution values of ATTR fibril maps shown in **A**.

### ATTR fibrils are exceptionally stable

Both ATTRwt and ATTRv fibrils proved extremely stable (Figure 5). The estimated free energy of these fibrils per chain ranges from −48.6 and −67.3 kcal/mol (Figure 5A). We compared their thermodynamic stability to the amyloid structures reported in the Amyloid Atlas by January 2022 and found that ATTR fibrils are significantly more stable than other fibrils, both by residue and chain (Figure 5B)(*13*, *14*). Analysis by residue reveals that the stability of ATTR fibrils results from the contribution of three pockets—the inner interface of the hairpin formed between Val 14 and Val 32, the inner pocket of the arch formed between residues Trp 79 and Phe 95, and the triquetra that connects the C and N terminal fragments together (Figure 5C). Most ionizable residues of ATTR fibrils are exposed to the outside, thereby enabling neutralization by other interactions that also contribute to the fibril stability. These include π-π stacking of aromatic residues (Figure 5D), hydrogen bonding from the stacking of asparagines along the fibril (Figure 5E), and hydrogen bonding of exposed ionizable residues within the same chain (Figure 5F). The loss of the C-terminal segment ending in Gly 67 in ATTRI84S fibrils from patient 1 may affect the overall stability of the fibril (−48.6 kcal/mol), although the stability per residue remains comparable to the other ATTR structures (−0.65 kcal/mol) (Figure 5A).

**Figure 5.**
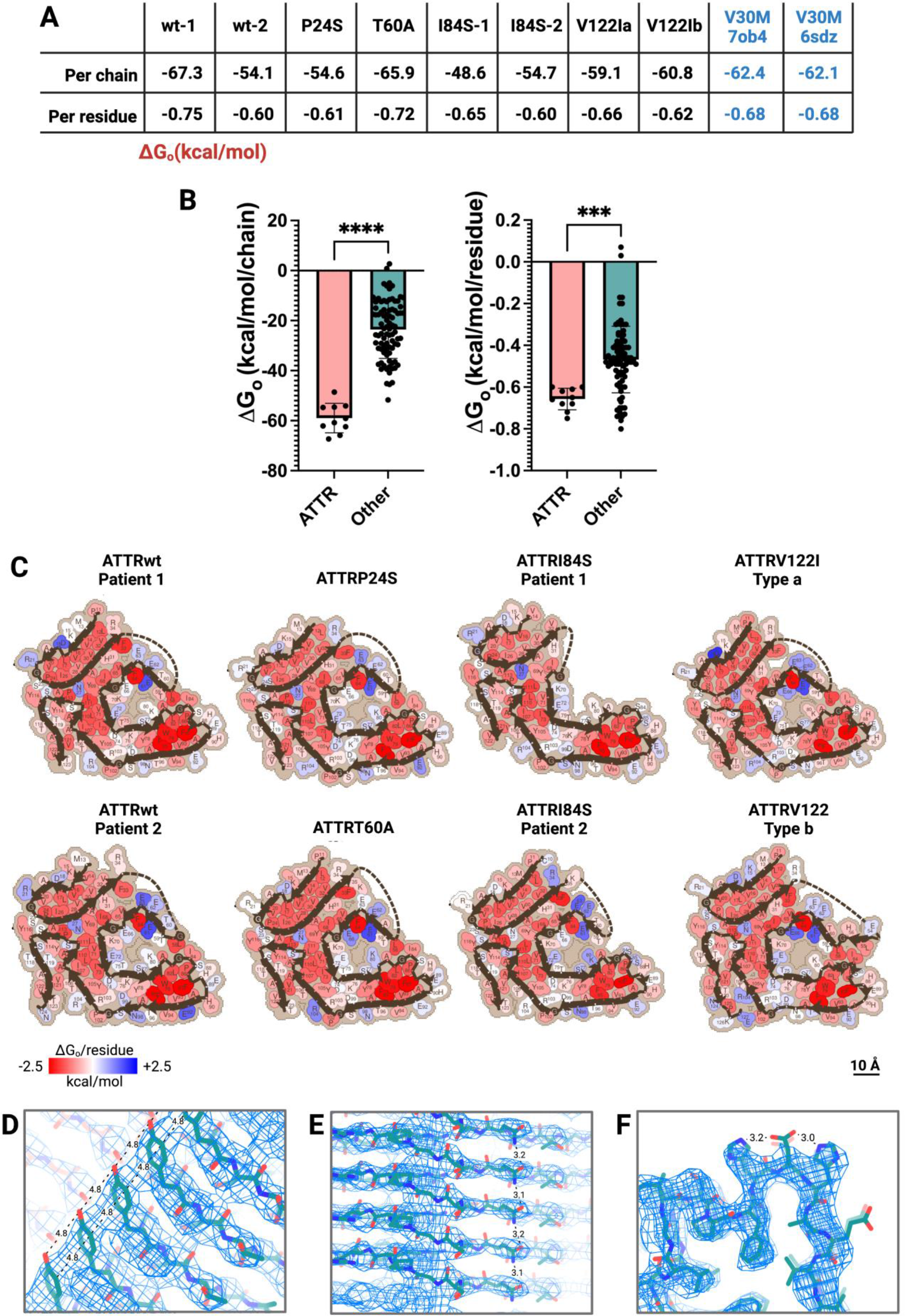
Fibril stability in ATTR amyloidosis. **(A)**. Stabilization energies per chain and per residue of ATTR fibrils in kcal/mol. (**B)**. Comparison of stabilization energies between ATTR fibrils and all amyloid fibrils reported in the Amyloid Atlas 2022(*13*). Left, stabilization energies by chain. Right, stabilization energies by residue. ATTR fibrils are significantly more stable than the other amyloid structures, considering the energies of both chains and residues. ***, p<0.001; ****, p<0.0001. **(C)**. Representation of stabilization energies per residue of cardiac ATTR fibrils determined in this study. Strongly stabilizing side chains are colored red, and destabilizing side chains are colored blue. **(D, E and F)**. Ionizable side chains on the outside of ATTR fibrils can be neutralized by π-π stacking **(D)**, hydrogen bonding stacking **(E)**, or intra-chain hydrogen bonding **(F)**. These three panels show examples found in ATTRwt fibrils from patient 1.

## Discussion

Our study reveals that cardiac fibrils from various patients of ATTR amyloidosis exhibit structural heterogeneity across genotypes, or within the same genotype of different patients, or within the same patient (Figure 6). These differences can be observed at the polar channel, at the ends of N or C terminal fragments, or in the fibril crossovers. While fibril structures of ATTRwt(s), ATTRP24S, ATTRT60A, and ATTRV122Ia contain the sequence from Leu58 to Gly67 to enclose the polar channel, in ATTRI84S patient 1, this entire sequence is missing, thus exposing the polar residues to the surrounding solvent. Interestingly, the two types of fibrils obtained from the ATTRV122I patient exhibit different helical twist angles (Figure 2), and display subtle changes in the folding, with additional interfaces formed by C-terminal amino acids in the fibrils with the faster twisting (Figure 2D-E). Finally, the structure of ATTRv fibrils exhibits local perturbations in resolution whereas the resolution of ATTRwt fibrils is homogeneously dispersed (Figure 4).

**Figure 6.**
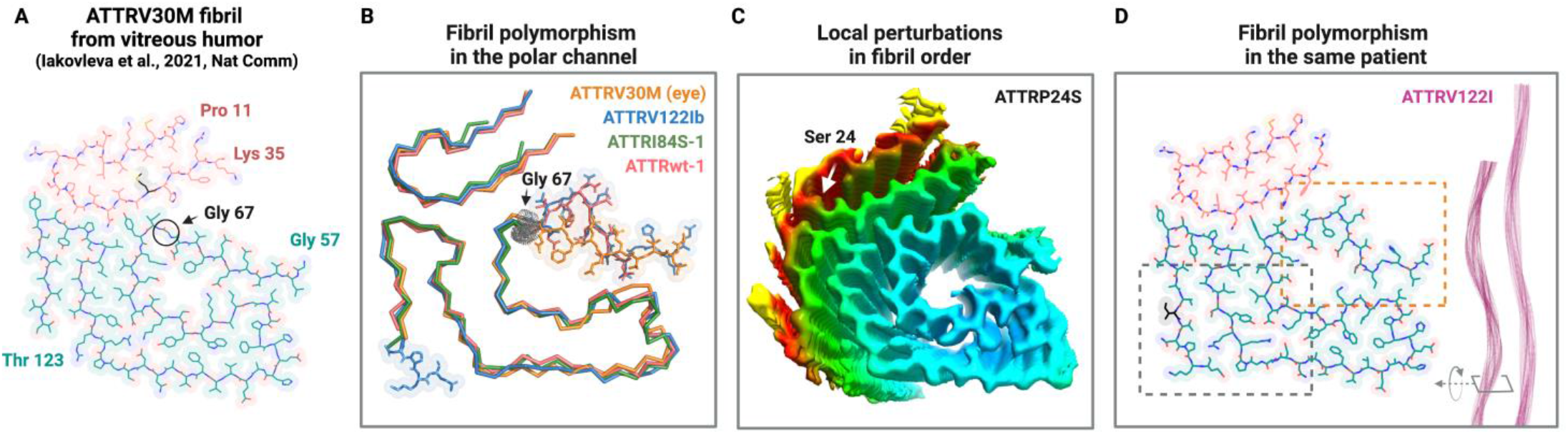
Fibril polymorphism in ATTR amyloidosis. **(A)**. Cryo-EM model reconstruction of one of the two protofilaments present in ATTRV30M fibrils from vitreous humor(*11*). The C terminal segment is colored pink. The N terminal segment is colored turquoise. The segment Leu 58 to Gly 67 crosses the polar channel diagonally, instead of in the shape of a pentagon. **(B, C, and D)**. ATTR fibril heterogeneity is structurally manifested at several levels. (**B)**. The segment that ends at Gly 67 folds differently in several ATTR fibril structures. For instance, in ATTRV30M fibrils from vitreous humor, this segment crosses the channel diagonally (blue). In the heart, this segment can be found surrounding the channel with the shape of a pentagon, such as in ATTRwt patients (red) or be disordered such as in ATTRI84S patient 1 (green). (**C)**. Some ATTR mutations, such as ATTRP24S shown in this panel, display local resolution perturbations that may be associated with structural disorder. **(D)**. ATTRV122I fibrils deposit in two conformations that differ in fibril twist and the presence or absence of additional interfaces (dashed square).

Our study shows that the structural heterogeneity in ATTR amyloidosis varies from patient to patient, instead of being specific to the disease (Figure 6). These results, therefore, challenge the current paradigm of “one disease equals one fibril polymorph” as proposed in tauopathies and synucleinopathies(*15*). Since all *ex-vivo* fibril structures determined to date depict wild-type sequences, with the exception of the two ATTR fibril structures described prior to our study(*8*, *15–21*), we expect that familial mutations of tau or α-synuclein, for instance, could lead to novel conformations.

The analysis of ATTR fibril stability reveals unexpected observations and informs us about transthyretin amyloidogenesis. Compared to other amyloid fibrils(*13*), ATTR fibrils exhibit the greatest stability estimated to date (Figure 5). In ATTR fibrils, the steric zipper motifs bring together three hydrophobic pockets that are the main contributors to the extraordinary stability of these fibrils. These are two β-arches (Val 14 to Val 32 and Trp 79 to Phe 95) and a hydrophobic triquetra that connects the C and N terminal fragments (Figure 5C). Many π-π interactions from the stacking of aromatic residues and hydrogen bonding from stacking of Asn 27, Asn 98, and Asn 123 along the fibril also contribute to the stability of the fibrils (Figure 5D, 5E). Exposure of most of the ionizable residues to the outside of the ATTR fibril or into the polar channel allows for the neutralization of their charges by posttranslational modifications, or hydrogen bonding with waters and ligands (Figure 5D-F). This extraordinary fibril stability likely results in greater resistance to proteolysis, denaturation, and clearance.

One observation from the stability estimates is that mutations in ATTRv cause structural perturbations in fibrils but do not affect the fibril stability when compared to ATTRwt, (Figure 4A & 5A). This suggests that the role of the mutations is more likely attributed to lowering the kinetic barrier to amyloid fibril formation, rather than stabilizing the fibril itself. This substantiates the notion that ATTRv mutations accelerate disease onset by decreasing the dissociation energy barrier of transthyretin tetramers, rather than promoting fibril formation(*22*, *23*)

A second observation is that the conformational variability in the polar channel, through segment Leu 58 to Gly 67, does not affect ATTR fibril stability significantly (Figure 5). In ATTRwt, ATTRP24S, ATTRT60A, and ATTRV122I fibrils of this work, and in previously published ATTRV30M from Schmidt *et al*, this segment surrounds and closes the polar channel in a pentagon-like shape that encompasses residues Leu 58 to Ile 84 (Figures 1, S3 and S4)(*10*). In ATTRV30M fibrils extracted from vitreous humor, however, this segment acquires an alternative fold that crosses the channel diagonally (Figure 6A)(*11*). In ATTRI84S fibrils from patient 1 but not from patient 2 this segment appears disordered (Figure 3). In all these cases, the pivotal residue Gly 67 governs the conformational polymorphism of ATTR fibrils, opening or closing the polar channel. And this polymorphism does not translate into significant changes in fibril stability (Figure 5), perhaps suggesting that the polar channel may not play an important role for amyloidogenesis and/or pathogenesis.

Several studies suggest a potential connection of ATTR amyloid fibril polymorphism with clinical presentation. Histopathological studies of ATTR fibrils reveal two distinct fibril types: type A, composed of full-length and fragmented transthyretin, and type B, composed exclusively of full-length transthyretin^24^. Type A and B fibrils are associated with differential clinical presentations in ATTRV30M amyloidosis patients^24^. We showed that the presence of fragmented transthyretin in ATTR fibril extracts has a positive correlation with their potential to seed further fibril formation^25^. Consistently, Type B ATTR patients have delayed cardiac deposition of wild-type transthyretin after liver transplantation than do Type A ATTR patients, who develop rapid cardiac deposition post-surgery^26,27^. All the samples used for cryo-EM structure reconstruction in this paper contain type A fibrils^25^. The structural variability that we found in ATTR fibrils indicates that type A pathology can consist of structural polymorphs, but with the limited information we have about the phenotypes of these patients, little can be said about the implications of our findings in pathology.

While more information is needed to fully grasp the implications of these findings, our studies highlight the high variability of fibril structures in this disease. Any correlation that exists between the disease phenotype and fibril polymorphs will require wider sampling of fibrils from other mutations and organs.

## Acknowledgments

In memory of Late Dr. Merrill D. Benson, who contributed greatly to the understanding of amyloid diseases and helped affected families for decades. We thank Justyna Kurleto, Morgan Schackmuth, and Drs. Romany Abskharon and Barbara Kluve-Beckerman for their technical support and useful discussion. We thank Aline McKensie for manuscript editing. Special thanks to the patients and families who generously donated tissues. We thank the UTSW Cryo-Electron Microscopy Facility, the UTSW Structural Biology Laboratory, the UTSW Electron Microscopy Core Facility, the UCLA California NanoSystems Institute, and the national cryo-EM facilities Stanford-SLAC (project CA60), PNCC (project 51267), and National Center for Cryo-EM Access and Training (NCCAT) for instrumentation, technical support, and/or data collection. Finally, we thank all the Twitter users that reconciled our scientific dilemma about the shape of our ATTR fibril ([Twitter Poll">Twitter Poll](https://twitter.com/SaelicesL/status/1530736126503358465?s=20&t=WZOxlz4g4wWqoEwjC9O2Jw).)

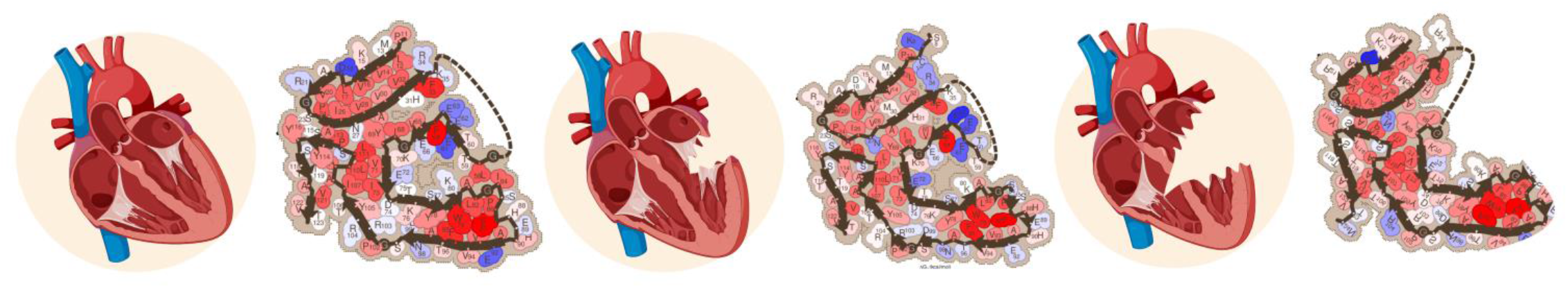

## Funding

American Heart Association (Career Development Award 847236)

National Institutes of Health, National Heart, Lung, and Blood Institute (New Innovator Award DP2-HL163810)

National Institutes for Aging, NIH grant R01 AG04812 Welch Foundation (Research Award I-2121-20220331)

UTSW Endowment (Distinguished Researcher Award from President’s Research Council and start-up funds)

Cryo-EM research was partially supported by the following grants:

National Institutes of Health grant U24GM129547, Department of Energy Office of Science User Facility sponsored by the Office of Biological and Environmental Research

Department of Energy, Laboratory Directed Research and Development program at SLAC National Accelerator Laboratory, under contract DE-AC02-76SF00515

NIH Common Fund Transformative High Resolution Cryo-Electron Microscopy program (U24 GM129539)

Simons Foundation (SF349247) and NY State Assembly and accessed through the NCCAT and the Simons Electron Microscopy Center located at the New York Structural Biology Center.

The Cryo-Electron Microscopy Facility and the Structural Biology Laboratory at UTSW are supported by a grant from the Cancer Prevention & Research Institute of Texas (RP170644).

The Electron Microscopy Core Facility at UTSW is supported by the National Institutes of Health (NIH) (1S10OD021685-01A1 and 1S10OD020103-01).

Part of the computational resources were provided by the BioHPC supercomputing facility located in the Lyda Hill Department of Bioinformatics at UTSW. URL: https://portal.biohpc.swmed.edu.

## Author contributions

Conceptualization: B.N., L.S.

Methodology: B.N., S.A., M.B., M.S., L.S.

Investigation: B.N., S.A., V.S., Y.A., R.P., M.C.F.R., M.S., P.S., Q.C., D.B., A.P., P.W., S.K., F.C.

Visualization: B.N., S.A., L.S.

Funding acquisition: D.S.E., L.S.

Project administration L.S.

Supervision: B.N., S.A., L.S.

Writing – original draft: L.S.

Writing – review & editing: B.N., S.A., V.S., Y.A., R.P., M.C.F.R., L.S.

## Data and Materials Availability

Structural data have been deposited into the Worldwide Protein Data Bank (wwPDB) and the Electron Microscopy Data Bank (EMDB) with the following EMD accession codes: 26587, 26685, 26688, 26691, 26692, 27323, 27324, 27325, and PDB accession codes: 8E7D, 8E7H, 8E7I, 8E7G, 8E7E, 8E7J, 8E7K, 8E7L. The PDB accession codes for the previously reported coordinates of ATTRV30M fibrils from vitreous humor and heart are 7ob4 and 6sdz, respectively. All data generated or analyzed during this study that support the findings are available within this published article and its supplementary data files.

## Materials and Methods

### Patients and tissue material

We obtained fresh frozen cardiac tissues from ATTR patients carrying wild-type TTR (*n* = 2) or TTR mutations (ATTRP24S, ATTRT60A, ATTRI84S, ATTRV122I,*n* = 5). Details of each sample included in the study are listed in Table S1. Specimens from the left ventricle of either explanted or autopsied hearts were obtained from the laboratory of Dr. Merrill D. Benson at the University of Indiana. The Office of the Human Research Protection Program granted exemption from Internal Review Board review because all specimens were anonymized.

### Extraction of amyloid fibrils from human cardiac tissue

*Ex-vivo* preparations of amyloid fibrils were obtained fromfresh-frozen human tissue as described earlier(*10*). Briefly, ~200 mg of frozen cardiac tissue per patient was thawed at room temperature and cut into small pieces with a scalpel. The minced tissue or ~100 mg of lyophilized fibrillar extract was suspended into 1 mL Tris-calcium buffer (20 mM Tris, 138 mM NaCl, 2 mM CaCl_2_, 0.1% NaN_3_, pH 8.0) and centrifuged for 5 min at 3100 × g and 4 °C. The pellet was washed in Tris-calcium buffer four additional times. After the washing, the pellet was resuspended in 1 mL of 5 mg/mL collagenase solution (collagenase was dissolved in Tris-calcium buffer) and incubated overnight at 37 °C, shaking at 400 rpm. The resuspension was centrifuged for 30 min at 3100 × g and 4 °C and the pellet was resuspended in 1 mL Tris–ethylenediaminetetraacetic acid (EDTA) buffer (20 mM Tris, 140 mM NaCl, 10 mM EDTA, 0.1% NaN_3_, pH 8.0). The suspension was centrifuged for 5 min at 3100 × g and 4 °C, and the washing step with Tris–EDTA was repeated nine additional times. All the supernatants were collected for further analysis, when needed. After the washing, the pellet was resuspended in 200 μL ice-cold water supplemented with 5-10 mM EDTA and centrifuged for 5 min at 3100 × g and 4 °C. This step released the amyloid fibrils from the pellet, which were collected in the supernatant. EDTA helped solubilize the fibrils. This extraction step was repeated five additional times. The material from the various patients was handled and analyzed separately.

### Negative-stained transmission electron microscopy

Amyloid fibril extraction was confirmed by transmission electron microscopy as described(*24*). Briefly, a 3 μL sample was spotted onto a freshly glow-discharged carbon-coated grid (Ted Pella), incubated for 2 min, and gently blotted onto a filter paper to remove the solution. The grid was negatively stained with 5 μL of 2% uranyl acetate for 2 min and gently blotted to remove the solution. Another 5 μl uranyl acetate was applied onto the grid and immediately removed. An FEI Tecnai 12 electron microscope at an accelerating voltage of 120 kV was used to examine the specimens.

### Cryo-EM sample preparation, data collection, and processing

Freshly extracted fibril samples were applied to glow-discharged Quantifoil R 1.2/1.3, 300 mesh, Cu grids, blotted with filter paper to remove excess sample, and plunged frozen into liquid ethane using a Vitrobot Mark IV (FEI). Cryo-EM samples were screened on either the Talos Arctica or Glacios at the Cryo-Electron Microscopy Facility (CEMF) at University of Texas Southwestern Medical Center (UTSW), and the final datasets were collected on a 300 kV Titan Krios microscope (FEI) at three different facilities: the CEMF, the Pacific Northwest Center for Cryo-EM (PNCC), and the Stanford-SLAC Cryo-EM Center (S^2^C^2^) (Table S2). Pixel size, frame rate, dose rate, final dose, and number of micrographs per sample are detailed in Table S2. Automated data collection was performed by SerialEM software package(*25*).

The raw movie frames were gain-corrected, aligned, motion-corrected and dose-weighted using RELION’s own implemented motion correction program(*26*). Contrast transfer function (CTF) estimation was performed using CTFFIND 4.1(*27*). All steps of helical reconstruction, three-dimensional (3D) refinement, and post-process were carried out using RELION 3.1(*28*).

All filaments were manually picked using EMAN2 e2helixboxer.py(*29*). Particles were first extracted using a box size of 1024, and 256 pixels with an inter-box distance of 10% of the box length. 2D classification of 1024-pixel particles was used to estimate the helical parameters. 2D classifications of 256-pixel particles were used to select suitable particles for further processing. Fibril helix is assumed to be left-handed. We performed 3D classifications with the average of ~30k to 40k particles per class to separate filament types using an elongated Gaussian blob as an initial reference. Particles potentially leading to the best reconstructed map were chosen for 3D auto-refinements. CTF refinements and Bayesian polishing were performed to obtain higher resolution. Final maps were post-processed using the recommended standard procedures in RELION. The final subset of selected particles was used for high-resolution gold-standard refinement as described previously(*30*). The final overall resolution estimate was evaluated based on the FSC at 0.143 threshold between two independently refined half-maps (Figure S7)(*31*).

### Model building

The refined maps were further sharpened using phenix.auto_sharpen at the resolution cutoff(*32*). Previously published model of ATTR-V30M (pdb code 6SDZ) was used as the template to build all near atomic resolution models. Mutations, rigid body fit zone, and real space refine zone were performed to obtain the resulting models using COOT(*33*). All the statistics are summarized in Table S 2.

### Stabilization energy calculation

The stabilization energy per residue was calculated by the sum of the products of the area buried for each atom and the corresponding atomic solvation parameters (Figure S5)(*14*, *34*). The overall energy was calculated by the sum of energies of all residues, and assorted colors were assigned to each residue, instead of each atom, in the solvation energy map.

### Recombinant protein expression and purification

Recombinant protein samples were prepared as described previously(*24*). Briefly, monomeric transthyretin (MTTR) was expressed in *Escherichia coli* and purified by affinity in a HisTrap column (GE Healthcare Life Science). Peak fractions were combined and further purified by size exclusion chromatography on a Superdex S75 prep grade column (GE Healthcare Life Science) in sodium phosphate–EDTA buffer (10 mM sodium phosphate pH 7.5, 100 mM KCl, and 1 mM EDTA). Peak fractions were pooled and stored at −20 °C.

### Amyloid Seeding Assays

Amyloid fibril extracts were used to seed the formation of new fibrils from recombinant MTTR as we described previously(*35*). Briefly, we further purified the extracts by treatment with 1% sodium dodecyl sulfate in NaP-EDTA buffer and centrifugation at 13,000 rpm for 10 min (rotor FA-24×2, Eppendorf). This purification process was repeated two times, and soluble fractions were discarded. The sample was washed with NaP-EDTA buffer three times by centrifugation and sonicated in cycles of 5 seg on/5 seg off for a total of 10 min at the minimum intensity (18%). The total protein content in the seed preparation was measured by the Pierce BCA Protein Assay Kit (Thermo Fisher Scientific). 2% (w/w) seeds were added to 0.5 mg/mL recombinant MTTR in a final volume of 200 μL containing 5 μM thioflavin T (ThT) and 1x PBS (pH 7.4). ThT fluorescence emission was measured at 482 nm with absorption at 440 nm in a FLUOstar Omega (BMG LabTech) microplate reader. Plates were incubated at 37 °C with cycles of 9 min shaking (700 rpm double orbital) and 1 min rest throughout the incubation. Measurements were taken every 10 min (bottom read) with a manual gain of 1,000. Figures show ThT signal in relative fluorescence units (RFU). Fibril formation was confirmed by transmission electron microscopy.

### Statistical Analysis

Statistical analysis of fibril stability was performed with Prism 9 for Mac (GraphPad Software) using an unpaired *t* test. All samples were included in the analysis and all measurements are displayed in the graphs.

---

**Figure. S1.**
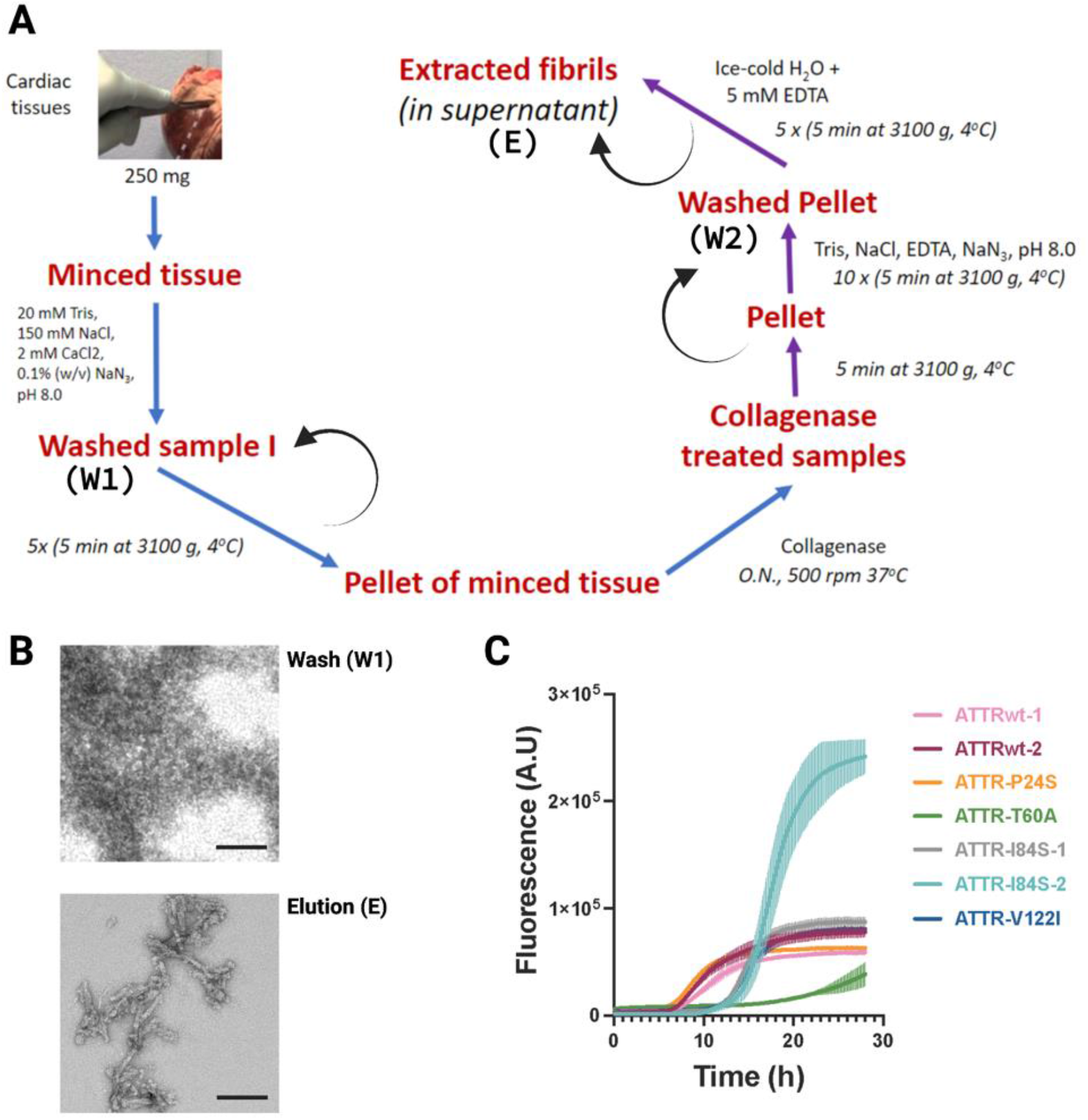
Extraction and assessment of ATTR fibrils from cardiac tissue. **(A)**. Schematic workflow for ATTR fibril extraction from human heart tissue. **(B)**. Representative negative stained TEM images of the first wash step (W1) of fibril extraction and the final fibril elution (E). Scale bar, 100 nm. **(C)**. ThT assay confirming that the fibrillar content of the elution can seed fibril formation of monomeric TTR at pH 7.4.

**Figure. S2.**
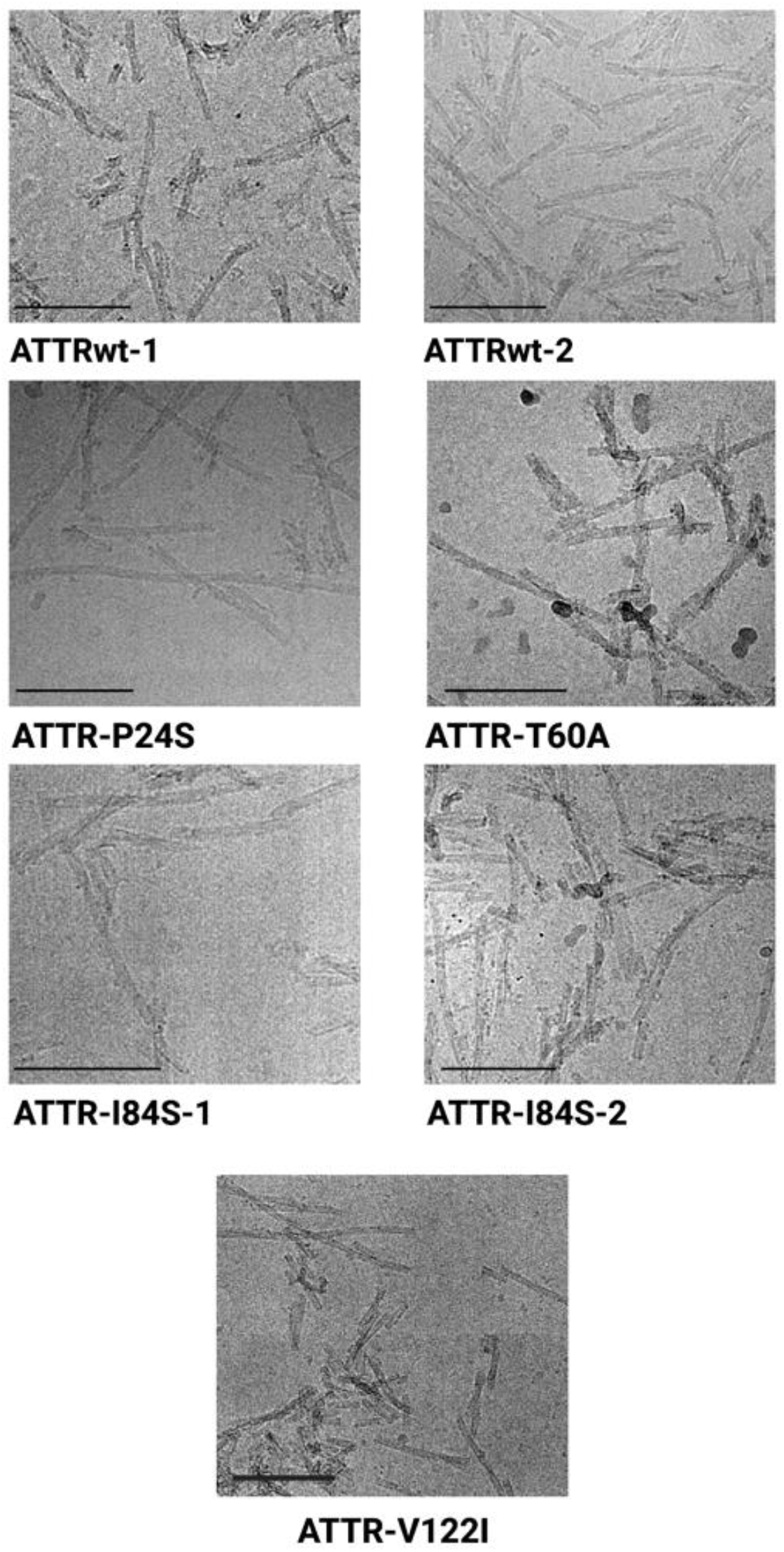
Representative cryo-EM micrographs of *ex vivo* cardiac fibrils. Scale bar, 100 nm.

**Figure. S3.**
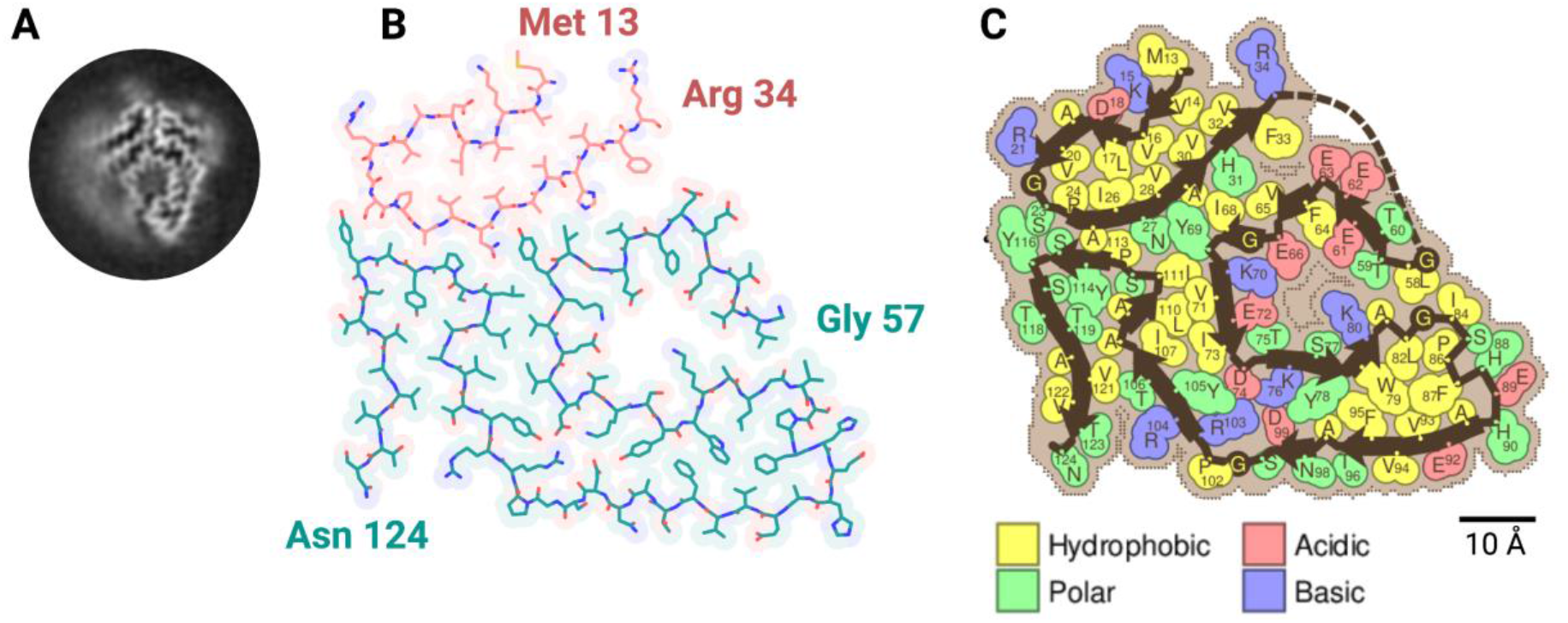
Cryo-EM structure of cardiac fibrils from ATTRwt patient 2. **(A)**. 3D class averages of curvy fibrils. **(B)**. Cryo-EM density and atomic model. Same as for the patient ATTRw-1, the model contains two fragments of transthyretin colored pink (residues Met 13 to Arg 34) and turquoise (residues Gly 57 to Asn 124). **(C)**. Schematic view of the fibril core showing residue composition.

**Figure. S4.**
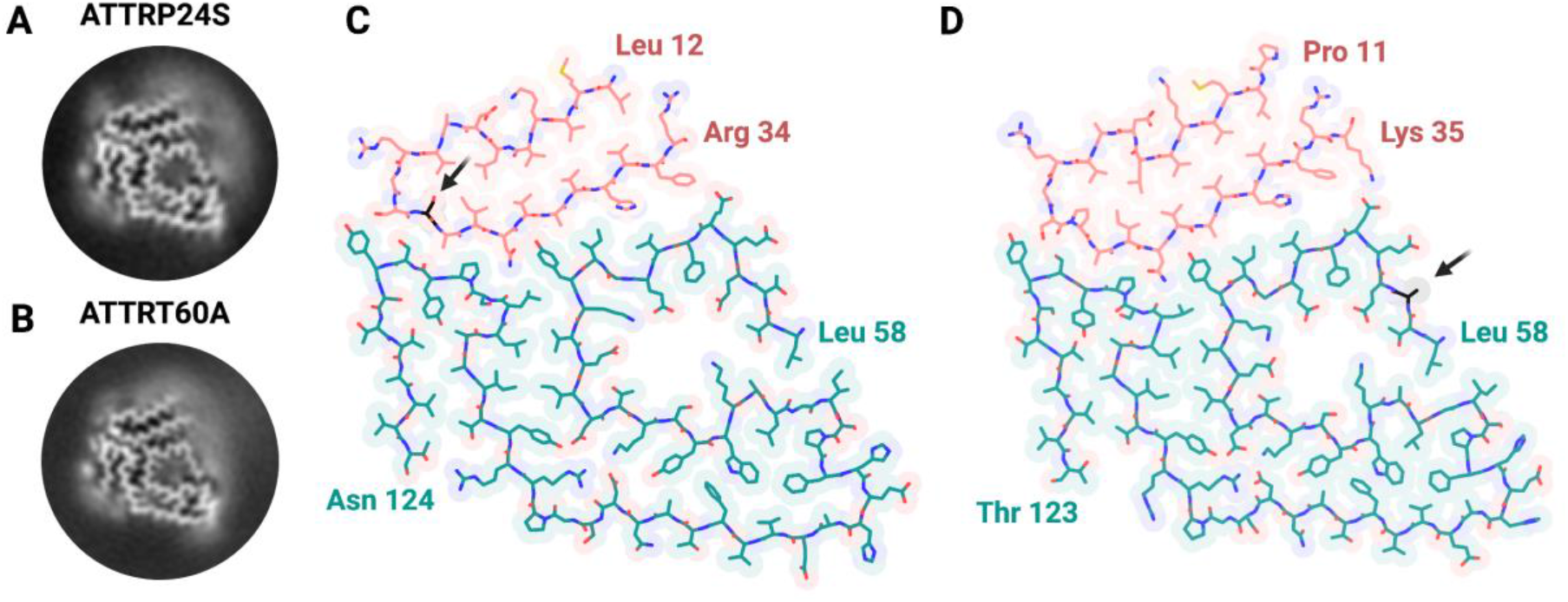
Cryo-EM structure of cardiac fibrils from ATTRP24S and ATTRT60A. **(A and B)**. 3D class averages of curvy fibrils from ATTRP24S (**A**) and ATTRT60A (**B**) patients. **(C and D)**. Cryo-EM density and atomic model of ATTRP24S (**C**) and ATTRT60A (**D**) fibrils. The N terminal segment is colored pink, encompassing residues Leu 12 to Arg 34 (**C**) and Pro 11 to Lys 35 (**D**). The C terminal segment is colored turquoise, encompassing residues Leu 58 to Asn 124 (**C**) and Leu 58 to Thr 123 (**D**). Arrows point at the mutation site.

**Figure. S5.**
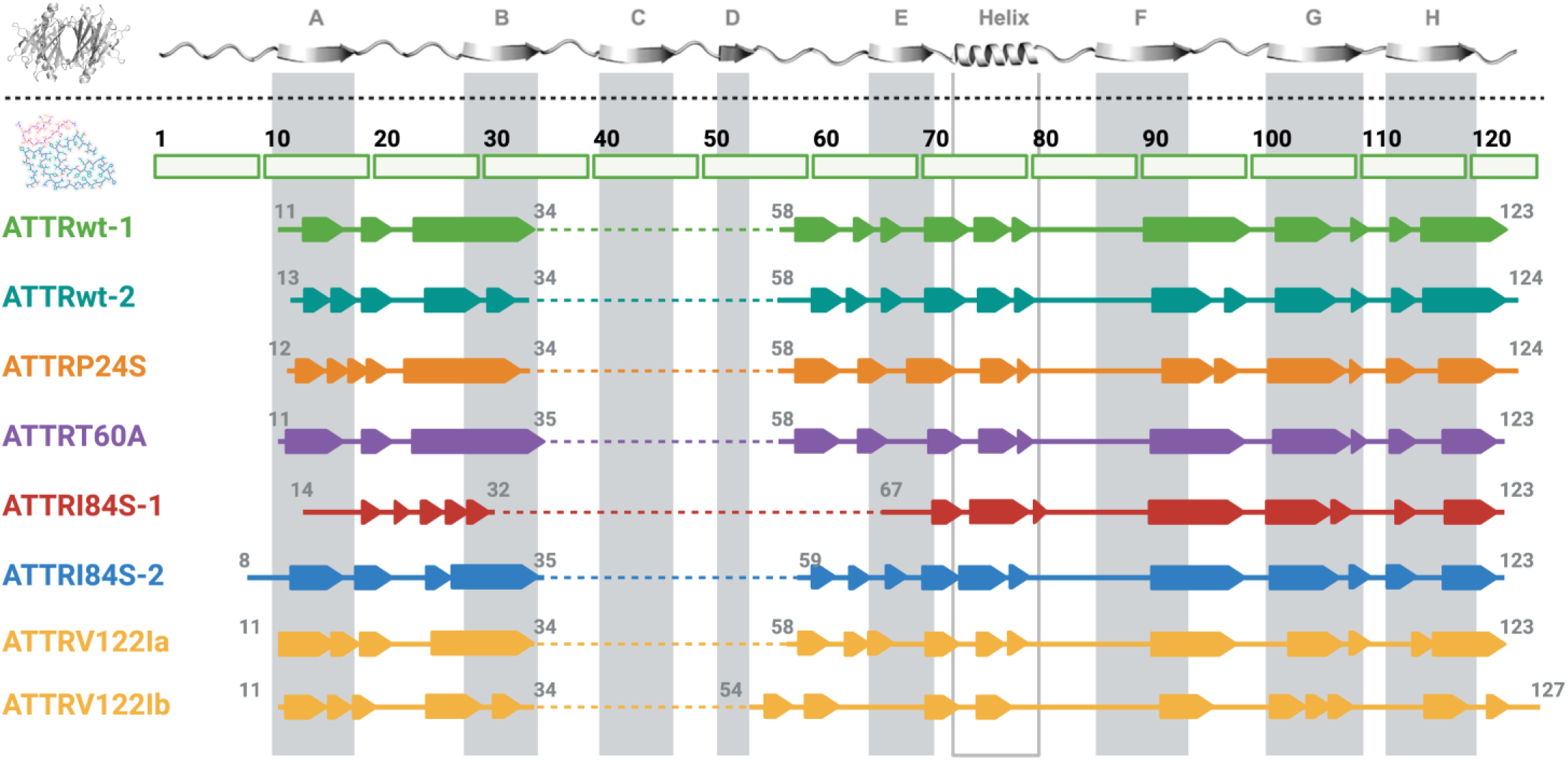
Secondary structure composition of ATTR fibrils. Schematic representation of the secondary structure of native transthyretin (above the dashed line) and fibrils from ATTRv and ATTRwt genotypes (below the dashed line). Fibrils are color-coded by patient.

**Figure. S6.**
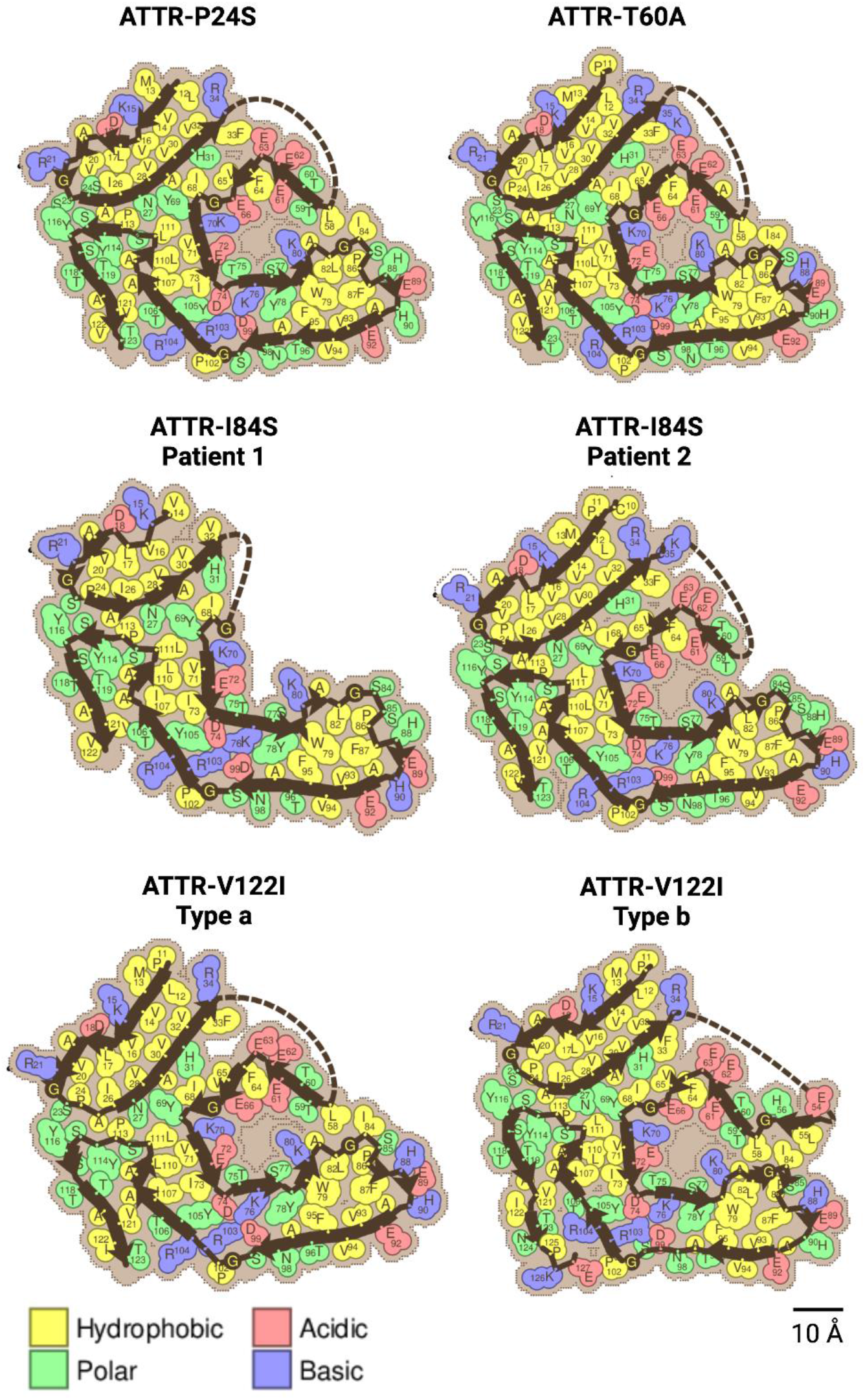
Schematic view of ATTRv fibrils showing residue composition. Residues are color-coded by amino acid category, as labeled.

**Figure. S7.**
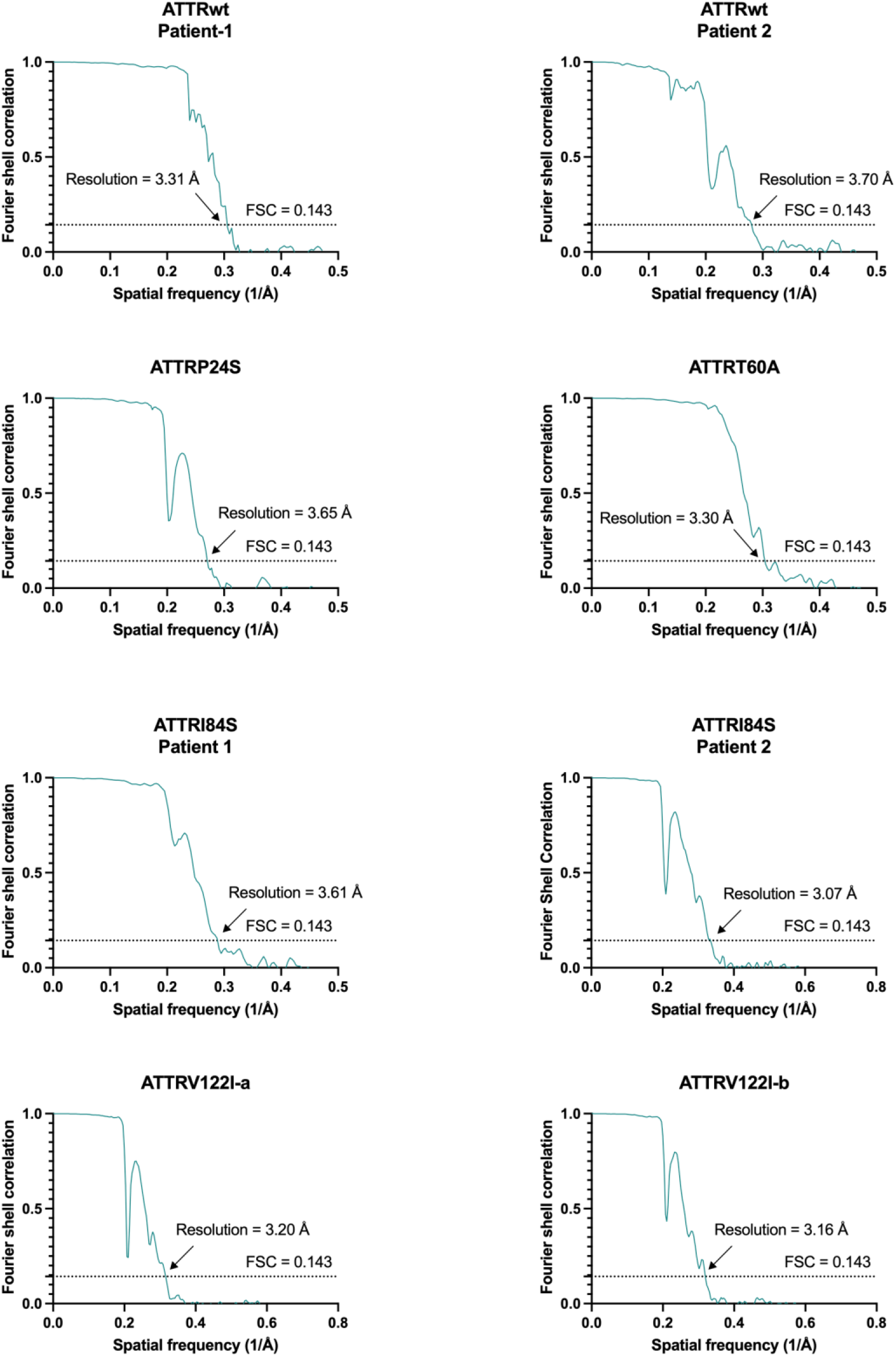
FSC curves between two half-maps.

## Supplementary Tables

**Table. S1.**
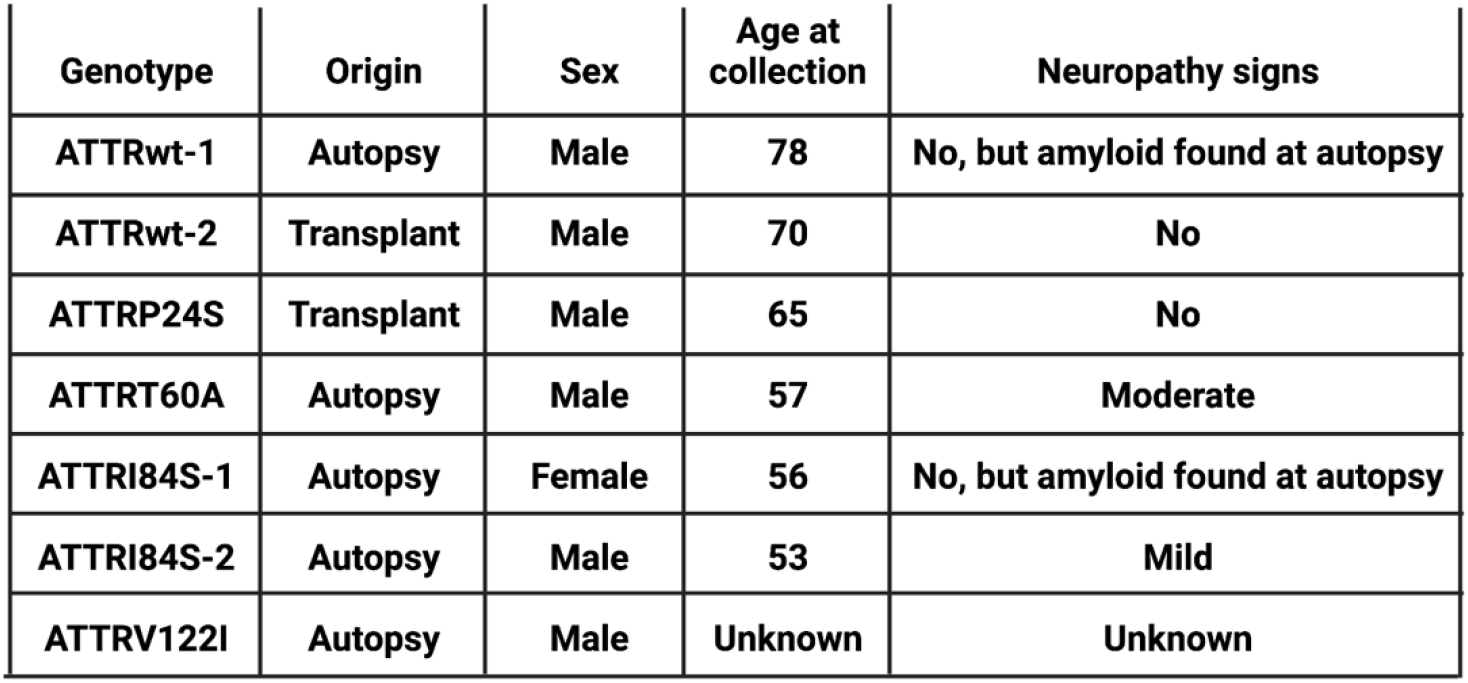
List of ATTR samples included in the study.

**Table. S2.**
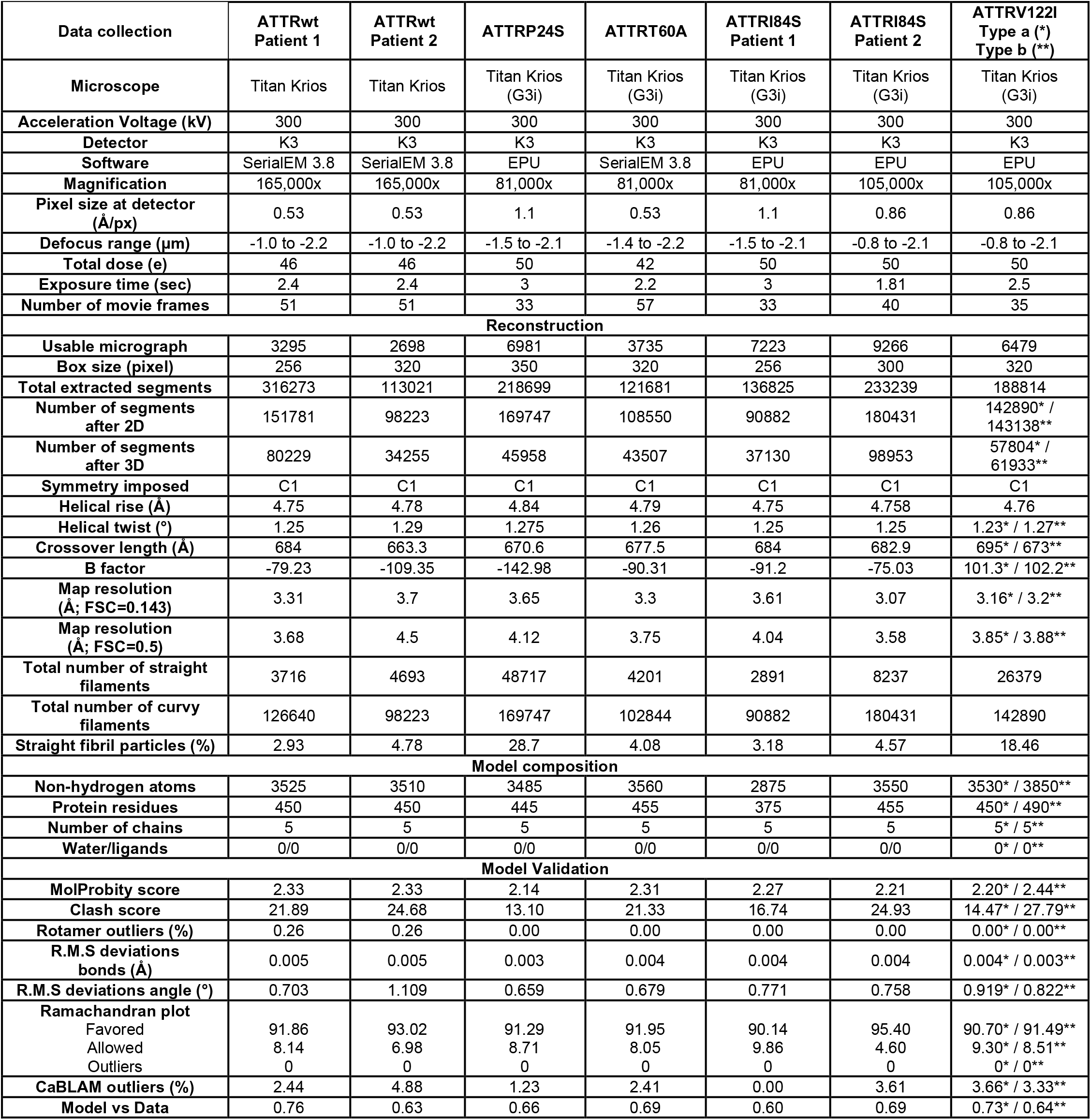
Data collection and refinement statistics.

